# A novel approach to digital characterisation of Tertiary Lymphoid Structures in colorectal cancer

**DOI:** 10.1101/2024.10.06.616904

**Authors:** Luis Munoz-Erazo, Saem Mul Park, Shelly Lin, Chun-Jen J. Chen, Lisa Y.Y. Zhou, Janet L Rhodes, Taesung Jeon, Sonya Fenton, John L. McCall, Roslyn A. Kemp, P. Rod Dunbar

## Abstract

Tertiary Lymphoid Structures (TLS) in cancer tissue are potential sites for the organisation of immune responses to cancer, and correlate positively with improved clinical outcomes for patients including in colorectal cancer (CRC). However it has proven challenging to standardise assessment of TLS due to the highly variable appearances of circumscribed domains of TLS within tissue sections. A recent three-dimensional reconstruction of TLS in CRC tissue showed that TLS are often large, multi-lobular structures, suggesting that assessing TLS across whole sections may be necessary to provide an accurate view of TLS activity in a patient’s tumour. In a pilot study we therefore used whole slide scans of multiplexed immunofluorescence images to characterise TLS from 22 subjects with CRC. Multiplexed staining for CD20, CD3, CD8, FoxP3 and Ki-67 enabled us to identify B-cells, CD8+ T cells, FoxP3– CD4 T-cells, and Foxp3+ CD4 T cells in all sections, and quantify both the presence of these cell subsets in lymphocytic clusters and their degree of proliferation within those clusters. In total we identified 524 lymphocytic clusters with morphology consistent with TLS. The count of TLS domains varied substantially between samples (from 4 to 100, mean 24) as did the proportion of total section area occupied by TLS (0.2%-7.8%) as well as size, morphology and cellular constituents; reflecting the intersection of the section plane with complex 3-dimensional structures. We quantified proliferation of B-cells and T-cell subsets within TLS domains across entire sections and compared to the canonical approach of counting and phenotyping individual TLS domains. The whole-slide approach proved far simpler than the canonical approach, generating digital summaries across our sample set clearly identifying patients with strikingly different levels of immune activity within their TLS. It also allowed us to demonstrate strong correlations between the proliferation of B-cells within CRC TLS and that of T-cell subsets. In summary we find that whole-section digital quantification of immune cell activity within TLS has major advantages over canonical approaches. Such whole-section approaches should accelerate research into correlations between TLS status and clinical outcomes, and ultimately enable standardised clinical tests based on automated analysis of multiplexed images.

## Introduction

Tertiary Lymphoid Structures (TLS) are structured organisations of immune cells that develop in non-lymphoid tissues following persistent pathogen infection, or in inflammatory conditions associated with autoimmune disorders, allograft rejection, and cancer (1, 2). TLS are closely related to other organised collections of lymphoid cells within normal tissues such as Peyer’s patches, and share many structural and organisational features with lymph nodes (LN). In sections of human tissue, TLS classically appear as circumscribed ovoid aggregates of lymphoid cells, centred around a B-cell follicle that may or may not contain a germinal centre (GC) comprising actively replicating B-cells. High endothelial venules (HEV) may be present within TLS, as well as peripheral T-cell zones with intercalating dendritic cells (1), though TLS lack other structural features of LN such as a capsule, with its connected hilum and trabeculae, or lymphatic sinuses.

In studies of cancer patients, correlations between TLS features and prognosis have been observed for several cancer types (1, 2) including colorectal cancer (CRC) (3, 4). The presence and characteristics of TLS in or around tumours have also been correlated with responses to immune therapy targeting the PD-1 pathway (5, 6), suggesting that TLS activity strongly influences the ability of T-cells to recognise and attack cancer cells.

Previous literature examining TLS features in cancer patients has typically sought to identify and characterise individual circumscribed zones within tissue sections that have features of TLS. Three classes of parameters have been extracted: the number and density of TLS within the section (7, 8); their location within the tumour with respect to cancer cells (9); and their cellular composition and functional state, especially their “maturity” (10). The manner in which all these parameters have been calculated and reported has been highly variable, making it challenging to compare TLS analysis in different clinical contexts.

Several previous studies set a threshold for the number of cells within a cluster needed to qualify as a TLS, though this threshold has varied (11–13). Other criteria for defining TLS have included cellular composition and architecture, such as the relative proportion of B-cells and T-cells within a cluster, and whether clear B- and T-cell zones are visible (4, 10, 11, 14, 15).

The location of TLS within tumours has also been variably reported. TLS are mostly found in peri-tumoural areas. In different human tumours, statistically significant correlations with prognosis have been found with both intra-tumoural TLS (16–18) and peri-tumoural/invasive margin TLS (10, 19). Hence, it remains unclear how TLS locations within tumours might impact prognosis.

Different criteria have been applied to classify the functional state of TLS. The presence of GCs and proliferating B-cells within B-cell follicles has often been used to describe TLS as “mature” and has been associated with improved prognosis (10, 20, 21). Some studies have also sought to sub-classify TLS according to the presence and density of particular cell types, such as follicular dendritic cells (FDC), CD8+ T-cells (22), and CD4+ T-cell subsets (23), including Foxp3+ CD4+ T-cells that are often regarded as “Tregs” (24), or the presence of HEVs (25). However, there is considerable heterogeneity in the cellular composition of TLS, even within a single tissue section (10, 23), and consensus has yet to emerge on how cellular composition might alter TLS function.

A large part of the variability in classifying TLS relates to the difficulty in assessing a dynamic three-dimensional (3D) structure from a single tissue section that effectively represents only a two-dimensional (2D) view. Recent 3D studies have begun to reveal human TLS as multi-lobular and interconnected (26) rather than discrete spheroidal entities. The appearance of TLS in a tissue section therefore depends on both the 3D shape of the TLS transected and the sectional plane, as illustrated in Figure 1. The sectional plane alters the shape and size of a circumscribed TLS domain in a tissue section (hereafter we use the term “TLS domain” to refer to this appearance in 2D). The sectional plane also changes the measurable cellular composition within a TLS domain, depending on whether it transects particular sub-structures such as GCs (Figure 1). Hence, the shape, size, and cellular composition of individual TLS domains within a tissue section are subject to the random interplay between the sectioning plane and TLS structures that might have highly variable multi-lobular shapes. This multi-lobular nature of human TLS (26) raises the possibility that instead of counting and classifying individual TLS domains, it might be preferable to characterise and summarise the totality of all these 2D cross-sections together.

**Figure 1:**
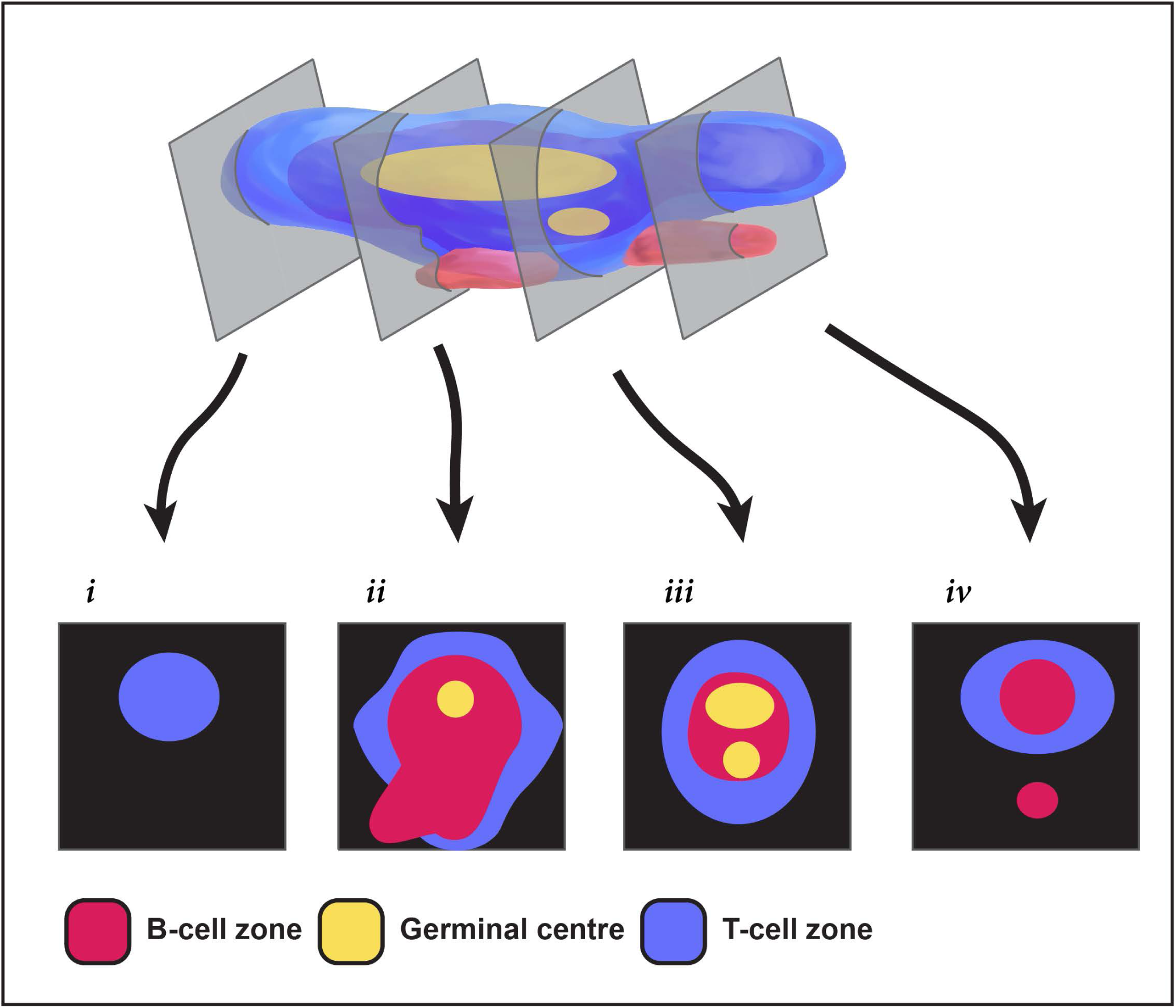
Challenges in extrapolating TLS structures from 2-dimensional sections. This pictorial representation shows how different a multi-GC TLS can appear in different 2-dimensional section planes. Section *i* shows only a T cell zone; sections *ii* & *iii* show differing relationships between T-cells, B-cells and the GC; section *iv* fails to capture a GC, and seems to show two TLS when the second is simply a protrusion from the same 3D structure.

We therefore used immunofluorescence (mIF) to image TLS across entire sections of tissue from 22 patients with colorectal cancer (CRC). As well as staining for B-cells and T-cell subsets, we stained for Ki-67 to quantify their proliferation, and identify GCs. We first used a canonical approach to quantification, counting individual TLS domains within each sample, and quantifying the cell subsets and their proliferation within each individual TLS domain. We then tested an alternative holistic approach: measuring the total area of TLS domains across the tissue section, and quantifying B- and T-cell parameters across this combined total area. This simple whole-section approach was far more efficient than the conventional method, yet still generated a similar ranking of patients for TLS activity. Our results suggest whole-section approaches to the digital characterisation of TLS will prove useful for comparative studies of large groups of cancer patients, and for future clinical applications.

## Results

### TLS domains in tissue sections from patients with CRC show highly diverse morphology

We examined formalin-fixed paraffin-embedded (FFPE) tissue sections from the tumours of 22 patients with CRC; patient characteristics are described in Table 2. We used the criteria shown in Methods to identify 524 TLS domains in these combined tissue sections (Figure S1).

TLS domains stained for the B-cell marker CD20, with variable staining for the T-cell markers CD3, CD8 and Foxp3 (Figure 2). Subsets of both B-cells and T-cells stained for Ki-67 (Figure 2). Most TLS domains were outside the tumour zones, with a minority in the peri-tumoural zones within 1mm from the invasive margin, and even fewer within the tumours; however, in this study we did not quantify location of TLS domains with respect to cancer cells.

**Figure 2:**
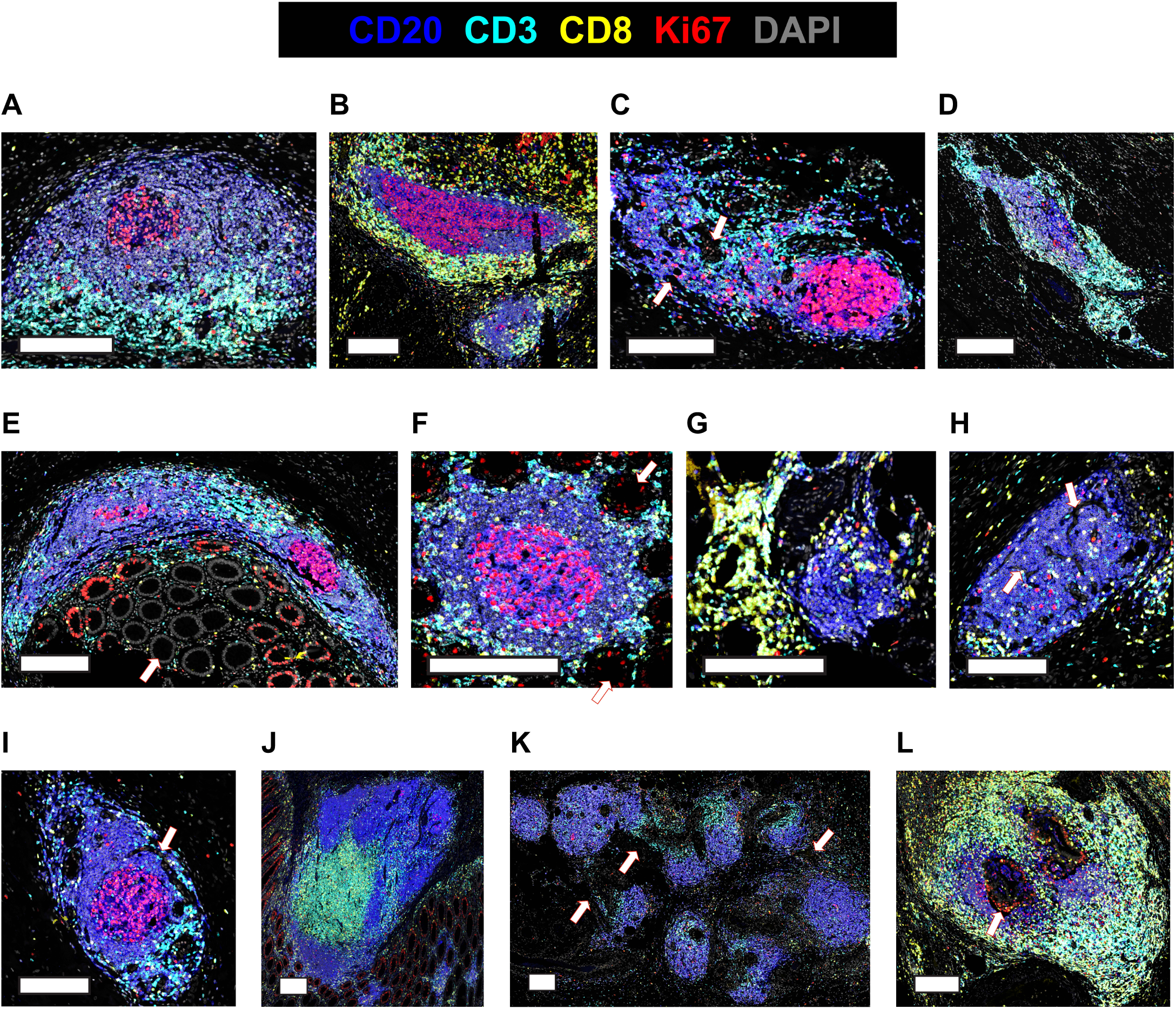
Diverse appearances of TLS in CRC. **(A-L)** Selected microscopic images of TLS imaged in the 22-patient cohort, highlighting the different visual appearances. Slides were stained for CD3, CD20, Ki67, CD8, DAPI and FoxP3 (not shown), imaged on the Akoya PhenoImager HT instrument. Scale bars are 200μm. Arrows are used to highlight **(C)** non-lymphocytic regions **(E,F)** epithelial crypts **(H,I)** channels **(K)** lymphocytic strands connecting discrete B-cell follicles **(L)** a Ki67+ tumour region encased by TLS.

In images of these TLS domains, some resembled the canonical descriptions of mature TLS (1), with a well-defined GC surrounded by a compact, mostly ovoid-shaped B-cell zone and an associated T-cell zone (Figure 2A). However, many TLS domains displayed other geometries. TLS domains were not always ovoid (Figure 2B, 2D) and both the B-cell zones (Figure 2B) and T-cell zones (Figure 2C) adjacent to GCs varied in shape, with non-lymphoid cells and ECM often intercalated (Figure 2C, arrows). Local tissue architecture also impacted TLS domain morphology (Figures 2D-G), while channels and holes sometimes led to perforated appearances, probably due to blood or lymphatic vasculature (Figure 2H-I).

Some TLS domains were large and contained multiple B-cell regions (Figure 2J), consistent with the section plane passing through large multi-lobular TLS (26). Other TLS domains appeared in close proximity, connected by strands of lymphocytes (Figure 2K, arrows), raising the possibility that they might be part of the same TLS in 3D. Occasional TLS domains surrounded Ki-67+ proliferating cells that were negative for CD3 and CD20, and therefore likely to be cancer cells (Figure 2L, highlighted by arrow).

Collectively, our results show that TLS domains do not usually appear as classical ovoid structures and often lack well defined T-cell zones within the sectioning plane, even when GCs are clearly present (Figures 2B, 2C, 2E, 2F, 2I). The spatial relationship of TLS to other tissue components and cancer infiltrates also affects the morphology of TLS domains visualised in 2D tissue sections.

### TLS domains show significant heterogeneity between and within CRC patients

We sought to quantify and classify individual TLS domains within each CRC tissue sample, by counting circumscribed TLS domains and phenotyping their cellular constituents (Figure S1). The counts of TLS domains within each tissue sample varied greatly (Figure 3A). The lowest and highest TLS domain counts were 4 and 100, respectively, with a mean of 24.

**Figure 3:**
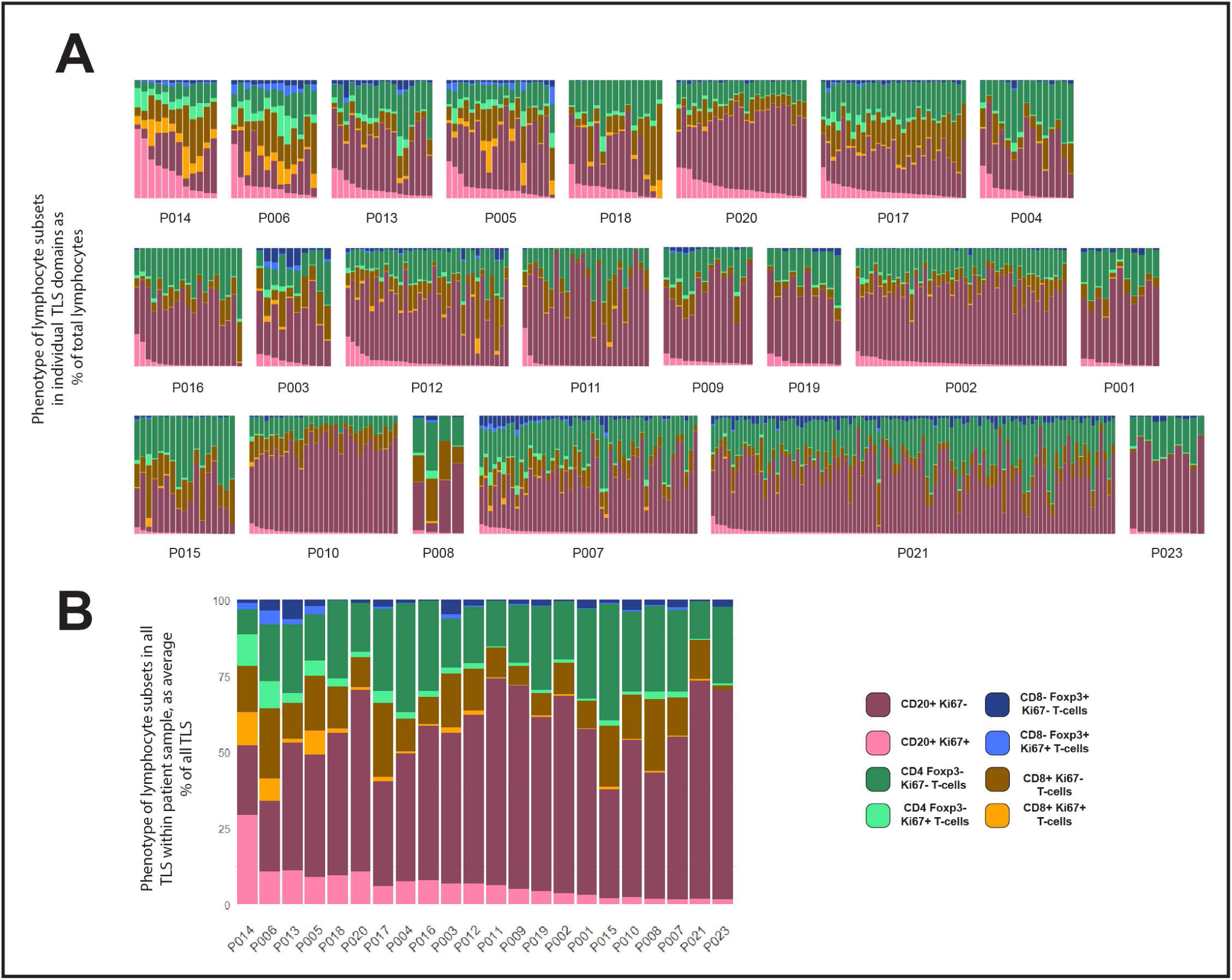
Variation within and between patients in lymphocyte composition of TLS domains. **(A-B)** 524 TLS domains across 22 patients with CRC were segmented and phenotyped, and lymphocyte subsets counted in each TLS. Data shown represent % of total lymphocytes occupied by each of the 4 lymphocyte types shown, with each lymphocyte subset sub-plotted for expression of Ki-67 (pale colours Ki-67+, dark colour Ki-67-). Patient identity codes are shown on the x-axis, and patients are ranked left to right according to average B-cell Ki-67+ % across all TLS within each patient. **(A)** Data from all individual TLS domains, grouped by patient. **(B)** Averages across all TLS domains in each patient.

The cellular composition of each TLS domain was analysed by digitally quantifying four different lymphocyte subsets, and the proportions of each of these subsets that expressed Ki-67 (Figure 3A). The proportion of B-cells expressing Ki-67 within TLS domains ranged from 0% to 94% (Figure 3A). Within each patient’s sample, the Ki-67+ proportion of B cells also varied considerably (Figure 3A). However, when comparing the results between patients across their TLS domains, differences between patients become apparent. Some patients had higher proportions of Ki-67+ B-cells in most B-cell zones (Figure 3A, upper rows), while others had consistently low proportions of Ki-67+ B-cells (Figure 3A, lower rows). Hence, despite the variation in Ki-67 expression by B-cells in different TLS domains within individual patient samples, patients with high B-cell proliferation within some TLS domains were readily distinguishable from those with very low levels of B-cell proliferation. Averaging the proportion of proliferating B-cells within all TLS domains in a tissue section allowed patients to be ranked for B-cell proliferation within their tissue sample (Figure 3B; this ranking was also used to order patients in Figure 3A).

Previous studies have suggested CRC patients with higher TLS numbers as well as those with a higher proportion of mature TLS have a better prognosis compared with patients with lower proportions (10, 23, 27). The numbers of TLS domains varied substantially even within the patients with the highest levels of B-cell activation (Figure 3B, top row). Additionally, patients among those with the lowest B-cell activation also had some of highest TLS domain counts (such as P007, P010 and P021). Taken together, these observations confirm that counts of TLS domains from 2D sections do not adequately summarise TLS activity.

### Ki-67+ status of B-cells is correlated with Ki-67+ status of T-cells, including Foxp3+ CD4+ T cells

Patients that ranked highest for the proportion of Ki-67+ B-cells also exhibited high Ki-67 expression in T-cell subsets (Figure 3B), suggesting a potential correlation between B-cell proliferation and that of certain T-cell subsets. We therefore compared the Ki-67 expression in B-cells with that in different T-cell subsets within individual TLS domains across all patients (Figure 4A). This analysis confirmed a correlation between the proportion of B-cells expressing Ki-67 within each TLS domain and the KI-67 expression of both Foxp3– CD4 T-cells and CD8+ T-cells (Figure 4A).

**Figure 4:**
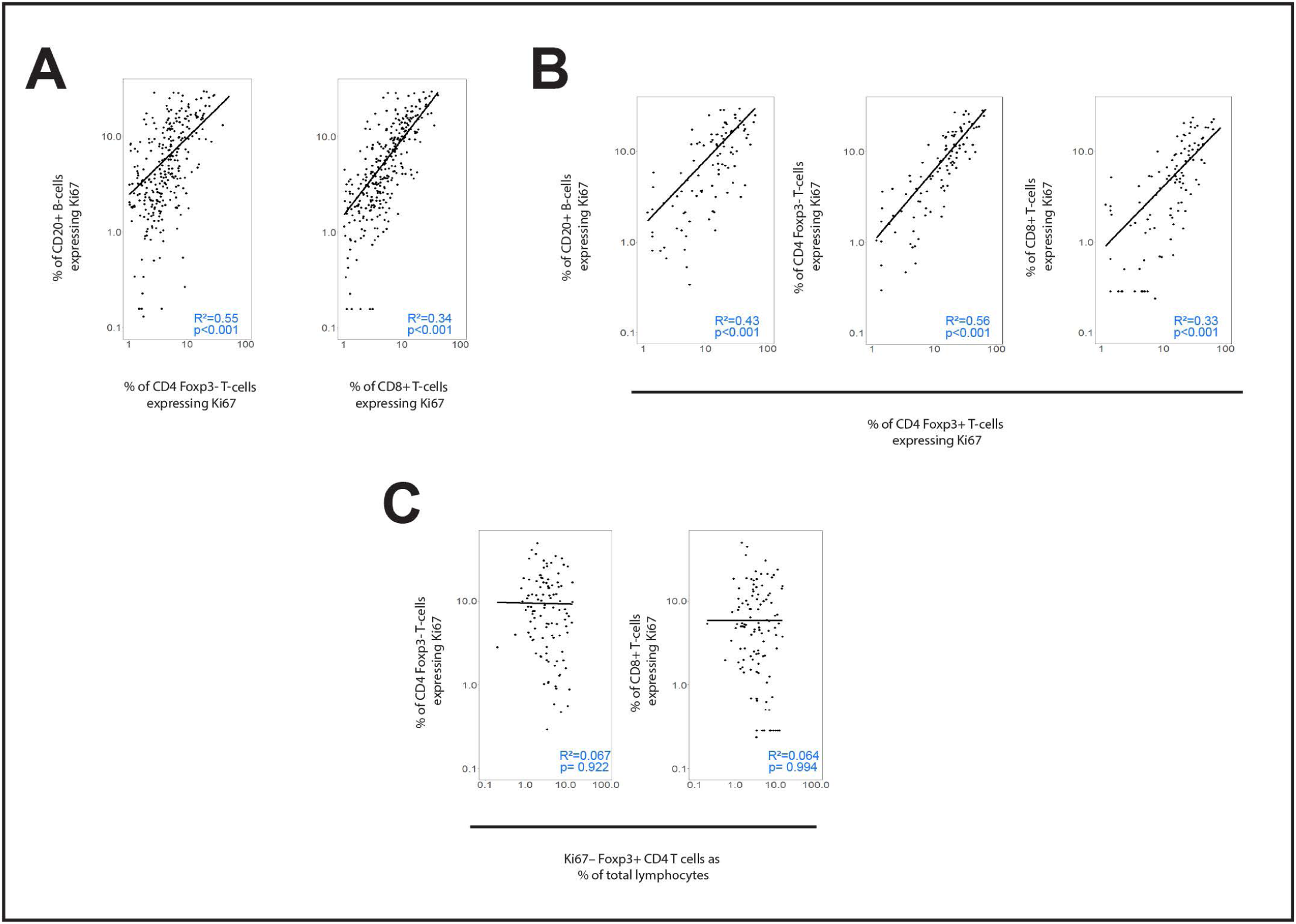
(A-C) Correlation of Ki-67 expression between lymphocyte subsets. Scatterplots showing the correlation between Ki-67 expression in different lymphocyte subsets within individual TLS domains. **(A)** Ki-67 expression in B-cells (y-axes) plotted against that in Foxp3– CD4 T-cells and CD8+ T-cells (x-axes) **(B)** Ki-67 expression in Foxp3+ CD4 T-cells (x-axes) plotted against that in B-cells, Foxp3-CD4 T-cells and CD8+ T-cells (y-axes) **(C)** Presence of Foxp3+ CD4 T-cells that are not proliferating (Ki-67–) in each TLS domain (x-axes) plotted against Ki-67 expression in Foxp3– CD4 T-cells and CD8+ T-cells. **(D-E) Automated identification of Germinal Centres (GC), and correlations with Ki-67 expression in B-cells and T-cells.** Individual TLS domains were classified digitally for the presence of GC, and TLS with (light colours) and without GC (dark colours) were plotted in each patient for the proportions of lymphocytes expressing Ki-67. **(D)** Proportions of B-cells expressing Ki-67 for TLS with and without GCs **(E)** Proportions of T-cells expressing Ki-67 for TLS with and without GCs.

Foxp3+ CD4 T cells also expressed Ki-67 frequently, and this expression correlated with that in B-cells, CD8+ T-cells and Foxp3– CD4 T cells (Figure 4B). Hence individual TLS domains with high proportions of Ki-67+ B-cells also had higher proportions of proliferation in all T-cell subpopulations, including Foxp3+ CD4 cells.

To test whether non-proliferating (Ki-67–) Foxp3+ CD4 T-cells might suppress T-cell proliferation, we plotted the correlation between the presence of this population (represented by their proportion of all lymphocytes within the TLS domain) and the Ki-67 expression in the other T-cell subsets (Figure 4C). The proportion of non-proliferating Foxp3+ CD4 cells per TLS domain did not show an inverse correlation with the presence of Ki-67 on other T-cell populations (nor B-cells) suggesting they were not suppressing T-cell proliferation within TLS.

### Assessing TLS area across entire tissue sections

Given the striking variation in the area of individual TLS domains (e.g. Figure 2), and the multi-lobular nature of TLS in at least some CRC samples (e.g. Figure 2K), it seemed likely that TLS count would poorly represent the total area of the TLS domains within a complete section. We therefore measured the total area of TLS within each section to compare this index of TLS volume with the canonical approach of counting the number of TLS domains.

Our data show that the count of TLS domains (Figure 5A) does not always reflect total TLS area within the section (Figure 5B). Plotting total TLS area in each tissue section against the count of TLS domains for that sample (Figure 5C) clearly shows the variability in the average area of individual TLS domains. For example, P006 and P013 are very similar in terms of TLS count (Figure 5A) but have very dissimilar total TLS area, reflecting the larger individual TLS domains in P013 than in P006, as is clearly evident in images from these subjects (Figure S3). Hence measuring total TLS area across an entire section better represents the total volume of TLS in that sample than counting individual TLS domains, because it accounts for the substantial variation in area of individual TLS domains.

**Figure 5:**
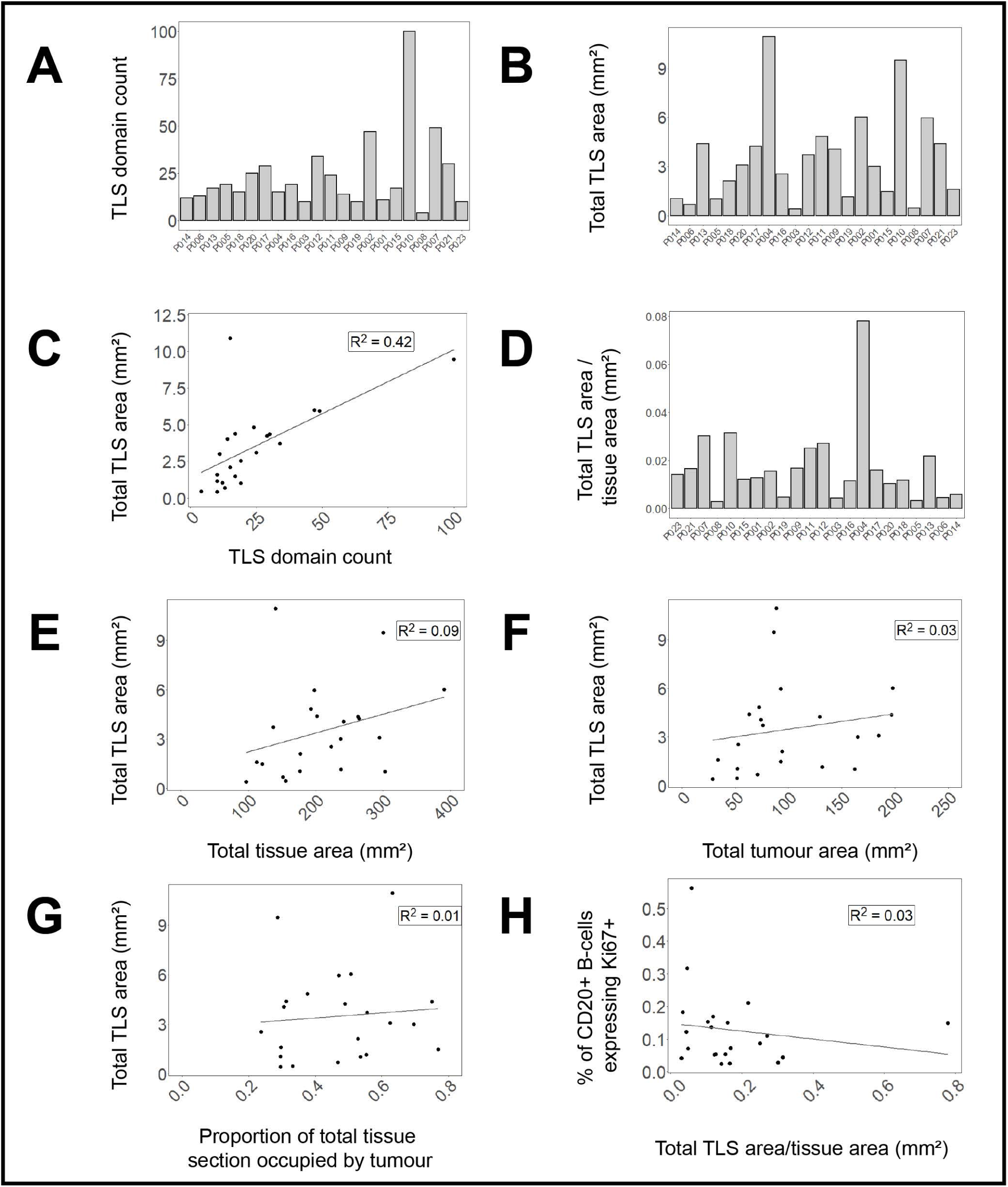
TLS domain density and correlation with other spatial measures. **(A)** Counts of circumscribed TLS domains in each tissue section **(B)** total area of TLS domains in each tissue section **(C)** average area of TLS domain per tissue section **(D)** proportion of total tissue area occupied by TLS domains. **(E-G**) Correlations of total TLS area across all tissue samples to **(E)** total tissue area **(F)** total tumour area **(G)** proportion of total tissue area occupied by tumour. **(H)** Correlation of B-cell Ki-67 expression (averaged across TLS domains within each sample) with the proportion of tissue section occupied by TLS in that tissue section.

Assessing total TLS area within a section needs to take account of total section size, and the proportion of the section that is occupied by tumour. The proportion of the total tissue area occupied by TLS varied substantially (Figure 5D) and there was no correlation between total section area and the area occupied by TLS (Figure 5E). Total TLS area did not correlate with total tumour area (Figure 5F) nor the proportion of the total section occupied by tumour (Figure 5G). Hence total TLS area varied independently of tissue section size and the degree of tumour infiltration.

To compare whether the volume of TLS within a section might correlate with TLS activation, we plotted the proportion of total section area occupied by TLS against the average proportion of B-cells expressing Ki-67 in each TLS domain (as shown in Figure 3B). No correlation was observed suggesting that the level of TLS activation represented by B-cell proliferation is not dependent on TLS volume.

### Assessing sum of B- and T-cell proliferation vs averaged individual TLS composition

Measurement of activation or proliferation of B- and T-cells across entire tissue sections may help correct for the failure of the section plane to transect structures such as GCs, while also simplifying analysis. We therefore re-analysed our data by combining data from all TLS domains identified within each section, and then ranking patients by the proportion of cell subsets across all TLS. When compared with the previous method based on identifying individual circumscribed TLS (Figure 3B-C), this whole-section approach led to broadly similar rankings of patient TLS activity (Figure 6A), with the same 8 subjects top-ranked, but with significant re-positioning of some patients due to the more holistic view of their B-cell Ki-67 expression (Figure S2). Plotting the proportions of Ki-67+ B-cells against Ki-67+ T-cells with this whole-section aggregated data showed improved correlations compared with the previous method based on individual TLS zones (Figures 6B compared with Figures 3A and 3B). This simpler whole-section analysis confirmed that proliferation of B-cells is more highly correlated with proliferation of both Foxp3– CD4 T-cells and CD8 T-cells than with proliferation of Foxp3+ CD4 T-cells (Figure 6B), and that proliferation of Foxp3+ CD4 T-cells was relatively rare compared with the other two T-cell subsets (Figure 6B).

**Figure 6:**
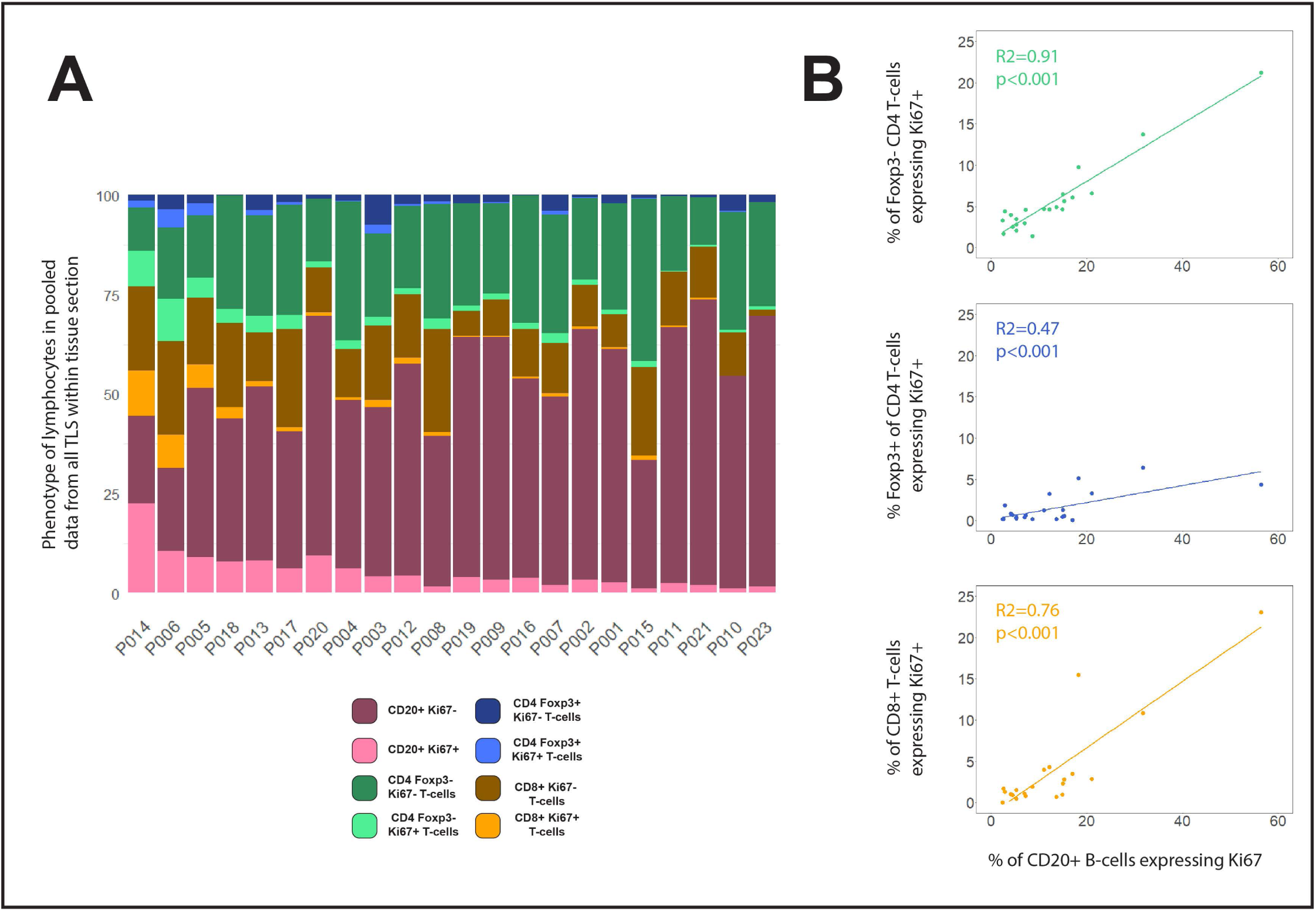
Compositions of total TLS area within each tissue section. **(A)** Lymphocyte phenotype data from all TLS domains within each tissue section were pooled before plotting % of each phenotype across the entire section. Patients were ranked by the proportion of B-cells expressing Ki-67. **(B)** Scatterplots showing the correlations between % of B-cells expressing Ki67+ with the Ki-67+ % for Foxp3– CD4 T-cells (green), Foxp3+ CD4 T-cells (blue) and CD8+ T-cells (orange). Spearman’s correlation test on all slopes were p<0.001.

## Discussion

The goal of this study was to create and evaluate a simple workflow to study TLS composition across entire sections of CRC tissue. We used mIF to stain CRC tissue, then digitally quantified the major lymphocyte populations in each TLS. We found the canonical approach of characterising individual TLS domains to be complex and difficult to summarise, given the highly variable appearance of TLS in a single section plane. We then developed a simpler method to quantify all TLS activity across entire sections of CRC, generating summary data for facile comparison between specimens. This whole-section approach generated a very similar ranking of patients for TLS activity to that generated by laboriously analysing individual TLS domains.

We stained for Ki-67 to identify proliferating lymphocytes. B-cells within GCs frequently expressed Ki-67, so the quantification of B-cell Ki-67 across whole tissue sections we performed might prove a much simpler measure of total GC formation within TLS than counting individual GCs. However we did note that in some TLS domains with high proportions of proliferating B-cells, the cells were not clustered into GCs (data not shown). Digital detection of B-cell clustering as well as B-cell proliferation might be necessary to accurately quantify GCs, though co-staining for other GC features, such as the presence of FDCs, may also prove useful in future.

We found Ki-67 staining to be particularly informative because it simultaneously reads out both B- and T-cell proliferation. Proliferation of all T-cell subsets within TLS correlated very well with B-cell proliferation when measured across entire sections, despite the highly variable T-cell content of individual TLS domains. This raises that possibility that quantifying both T-cell activation parameters as well as B-cell parameters within any structure that might be part of a TLS may add to predictive power for clinical outcomes; this approach contrasts with canonical analysis restricted to tissue neighbourhoods that contain both cell types juxtaposed.

Foxp3+ CD4 T cells are sometimes considered to be regulatory T-cells with potential to suppress activation and proliferation of other T-cell subsets. To check whether the non-proliferating Foxp3+ CD4 T cells might exert a negative effect on the proliferation of the other subsets, we tested the correlation between the presence of these cells in TLS and the Ki-67 signal in all T-cell subsets: no correlation – either positive or negative – was found. In contrast, we found a strong positive correlation between the proportions of Foxp3+ CD4 T-cells expressing Ki-67, and the proliferation of both CD4 and CD8+ T-cells, suggesting Foxp3+ CD4 T-cells had often been activated to proliferate within TLS alongside Foxp3– CD4 T-cells and CD8+ T-cells. These data suggest that the presence of Foxp3+ CD4 T cells within CRC tissue does not suppress proliferation of other T-cell subsets. This result might seem inconsistent with the immunosuppressive activity ascribed to Foxp3+ CD4 T-cells in some human cancers, where their presence within tumours has been associated with poor outcomes (29, 30). However while Foxp3 is often used as a marker of Treg cells, in humans, many Foxp3+ T-cells are not Tregs (31, 32) and may even be activated effector/memory cells (33, 34). Indeed the presence of CD4+ CD45RA– effector/memory T-cells expressing low levels of Foxp3+ has been associated with improved prognosis in CRC (32). Hence our data reinforce the concept that the expression of Foxp3 by CD4 T-cells does not necessarily correlate with immunosuppressive activity.

Our study suggests measures of total TLS area are likely to better represent the volume of TLS within a patient sample than counts of individual TLS domains. The average area of individual TLS domains varied substantially between patients, so that the number of discrete TLS domains did not accurately reflect the proportion of the total section occupied by TLS. Our study also showed no correlation between this TLS area and the presence of proliferation lymphocytes within the TLS, suggesting TLS volume and activation are independent. In future clinical studies it would therefore be valuable to report TLS area parameters separately from activation parameters in order to tease out correlations with clinical features.

We can already envisage several potential improvements to whole slide analysis of TLS that are worthy of further investigation:

Firstly we suggest that whole-slide approaches incorporating cancer cell markers might give a better index of overall proximity of TLS to cancer cells than merely marking tumour boundaries in 2D, given the complexity of the 3D spatial relationship between TLS and tumours. We did not formally assess the spatial relationships of TLS to cancer cells, but in addition to occasional intra-tumoural TLS we did note some TLS domains wrapped around isolated tumour buds. Such TLS may be functionally similar to intra-tumoural TLS given the direct access TLS-resident immune cells will have to cancer cell macromolecules (14). It may therefore be useful to quantify the area of TLS that is in contact with any cancer cells, no matter where they are located with respect to the main tumour body.

Secondly we recognise that further investigation of the constituents of immune cell clusters within CRC tissue may enable new classifications of clusters that could be independently quantified across entire sections. In this study we took a relatively broad view of immune cell clusters that might comprise parts of TLS within the tissue section, consistent with the 3D structure of human TLS (26). However use of additional markers that can distinguish other cell types, such as myeloid cell subsets, might reveal distinctive subtypes of clusters. We did not attempt to distinguish between TLS and other gut-associated lymphoid tissue (GALT), in the absence of markers that can distinguish pre-existing GALT structures from induced TLS (28). Should such specific markers emerge it would be useful to include these in future whole slide analysis.

In conclusion, we have shown that whole section analysis of TLS activity in aggregate efficiently averages the highly variable appearance of complex TLS structures within a single 2D section plane. Quantification of molecular markers such as Ki-67 in TLS lymphocytes across entire sections greatly reduced the complexity of canonical approaches based on counting and characterising individual TLS domains, and may potentially improve accuracy. These novel methods can now be tested against clinical parameters in large cohorts of patients, potentially with automated detection and aggregation of TLS domains, and inclusion of cancer cell markers to enable quantification of TLS proximity to cancer cell zones.

## Materials and methods

### Patient selection

FFPE tissue blocks were obtained from 22 patients from the Dunedin Colorectal Cohort under ethical approval from the NZ Health and Disability Ethics Committee (HDEC 14/NTA/033). Previous H&E staining of these tissue blocks had revealed at least 3 highly-defined immune aggregates in each sample. Patient characteristics are explained in Table 1.

**Table 1:**
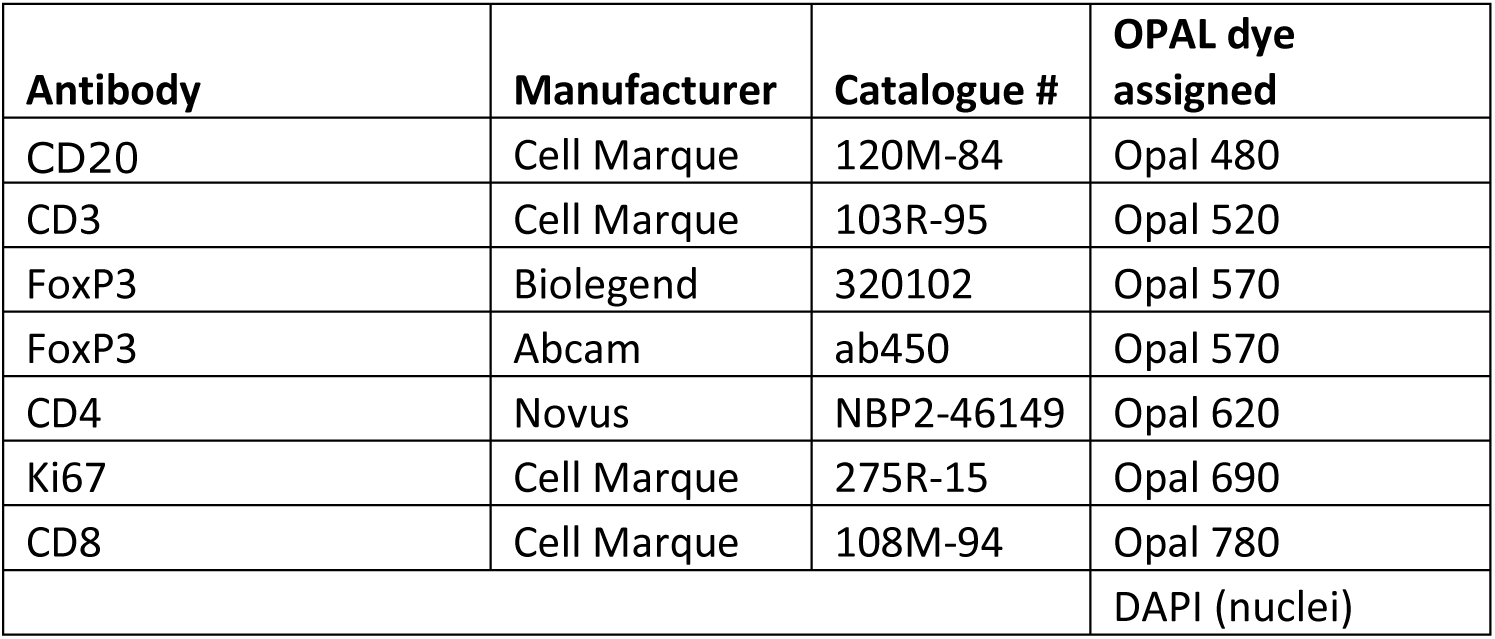
List of antibodies used in this experiment and the Opal dyes assigned to them. The order of the antibodies listed is the order that the antibodies were added used in the cyclical staining procedure Opal procedure.

**Table 2:**
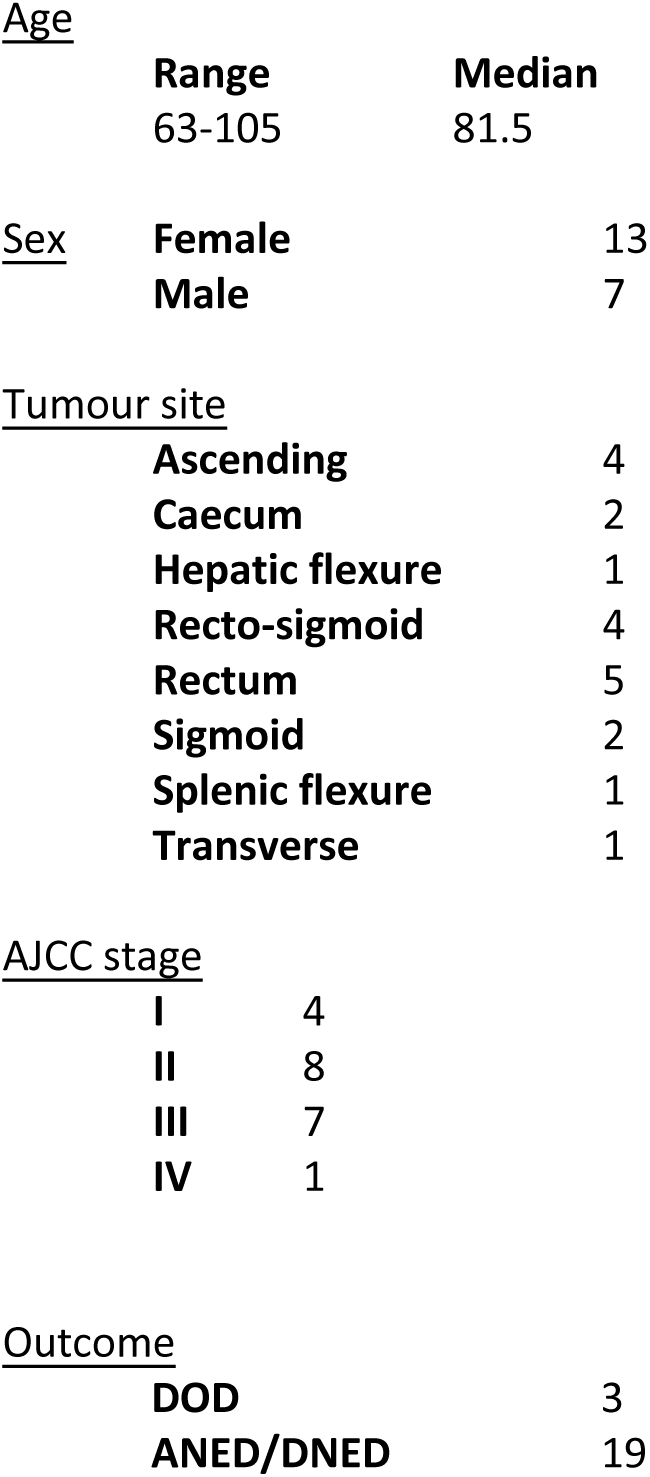
List of patient characteristics. AJCC= American Joint Committee on Cancer, DOD= Dead of disease, DNED=Died, No Evidence of Disease, ANED= Alive, No Evidence of Disease

### Tissue processing and imaging

5µm sections were derived from these blocks by the Histology Services Unit at the University of Otago. Sections were deparaffinised, antigen retrieved (AR6 solution, Akoya Bioscience) and blocked with 10% human serum (ThermoFisher). Slides were stained for markers using the 7-plex Opal Polaris kit (Akoya Biosciences) according to the manufacturer’s instructions, using the primary antibodies and Opal dyes listed in Table 1. Slides were mounted and imaged with the PhenoImager HT instrument (Akoya Biosciences). Due to the variable staining CD4 stain across the patient dataset, it was omitted from subsequent analysis. Hence our analysis utilised 5 markers in addition to the nuclear stain DAPI: CD3, CD8, Foxp3, CD20, Ki-67.

For H&E images, slides were deparaffinised and stained with Gills hematoxylin (Merck/Sigma Aldrich) counterstained with Eosin Y, aqueous (Merck/Sigma Aldrich) before dehydration and mounting with Eukitt Quick-hardening mounting medium (Merck/Sigma Aldrich).

### Image analysis

Image unmixing, cell segmentation and positive channel marker thresholding were undertaken using the InForm (version 2.6 AKOYA BioSciences) program. Phenotyping and analysis of InForm-derived data was undertaken using R (version 4.0.4) in conjunction with custom analysis pipelines and the phenoptr package (version 0.3.2). Additional area measurements and assessments using the fluorescence wholeslide scans were undertaken with QuPath (version, 0.4.2) guided by the equivalent H&E scans annotated by a pathologist.

To improve comprehension of our results, we classified CD3+ CD8-cells as CD4 T-cells, since almost all T-cells present in CRC are single positive for either CD8 or CD4 (35) (36). Lymphocytes were therefore classified into 4 groups: B cells (CD20+ CD3–); CD8 T-cells (CD3+ CD8+); Foxp3– CD4 cells (CD3+ CD8– Foxp3–); Foxp3+ CD4 cells (CD3+ CD8– Foxp3+). Each of these 4 cell subsets were then sub-classified as Ki-67+ or Ki-67– to generate counts of 8 cell classifications for each TLS. A ninth category of cells shown in some analyses is “undefined” – cells that did not fall into the aforementioned categories, the vast majority of which were negative for all markers.

### TLS quantification

Our strategy to determine TLS from other non-TLS lymphoid aggregates for downstream analyses is outlined in Figure S1. Lymphoid aggregates were initially identified in images by two researchers; where B-cell aggregates were close to each other and were connected by strands of lymphocytes (e.g Figure 2J) these were considered as single aggregate, The two researchers then re-assessed each other’s selections to determine which aggregates met defined criteria for classification as TLS domains. Cellular composition of each aggregate was phenotyped as described above, and only aggregates of ≥250 cells comprising >50% B-cells and/or T-cells were included as TLS domains.

Given the lack of consistency in the literature in how the peri-tumoural region is defined (1), TLS domains were not further sub-classified according to their distance from tumour margins, although all TLS domains analysed were either intra-tumoural or within ≈10mm of cancer cells.

### Area measurement

Total tissue area measurements were made on Qupath (37) using the pixel classifier thresholder function. Briefly, the average of all channels were prefiltered (Gaussian) and a threshold was chosen that coved the tissue region. Tumour area measurements were made manually, aided by H&E stained sections from the same tumour annotated by a pathologist (T. Jeon). TLS tissue measurements used the pixel classifier thresholder function using CD3 and CD20 channels. Manual adjustments were made to the region where required.

### Statistical analyses

Statistical analysis on individual TLS data was performed in R using the R package lme4 (v1.1-29) to apply a generalised linear mixed model with REML/residual maximum likelihood to account for grouping effects. The p-values were calculated with lmerTest (v3.1-3) using Satterthawite’s method. For the whole TLS tissue analysis, a linear regression with Spearmann’s correlation test was implemented using the stats package on R (v4.0.4). Values scaled in log10.

To reduce the effect of skew from small numbers of cells, TLS with less than 50 cells of the assessed phenotypes were removed from analysis when performing correlation analyses. This was also performed for the comparisons of GC-detected vs GC-not detected TLS.

---

**Figure S1:**
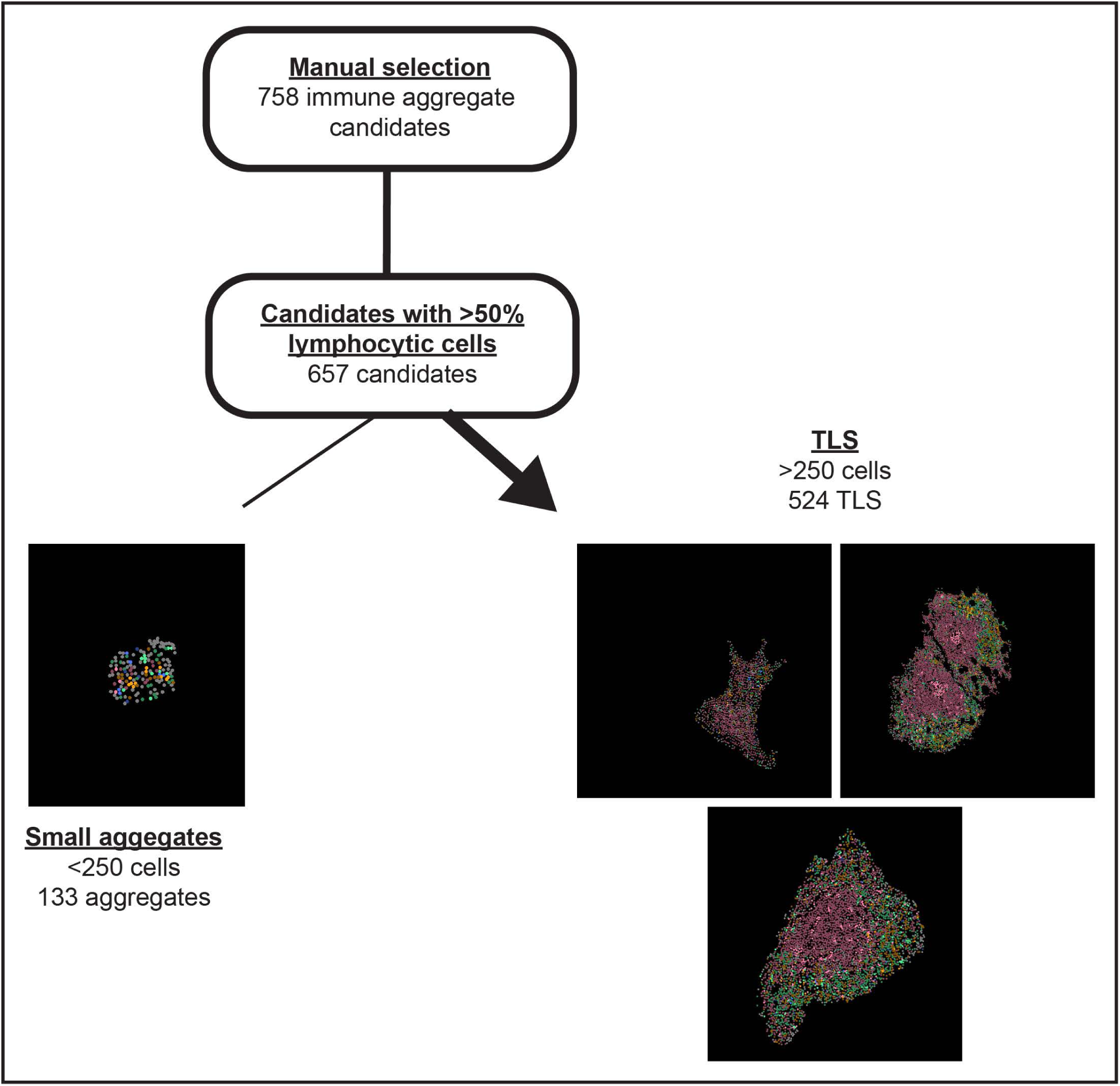
Flowchart of TLS analysis. All lymphocyte clusters with potential to be part of TLS were selected within regions of interest for cell segmentation, phenotyping and digital analysis, as described in the Methods. Clusters comprising ≥50% non-lymphocytic cells (“undefined” cells), were first removed from further analysis, followed by clusters comprising less than 250 cells. The 524 TLS domains that met these thresholds were further sub-classified as containing germinal centres (GC) or not, using the computational approaches described in the Methods.

**Figure S2:**
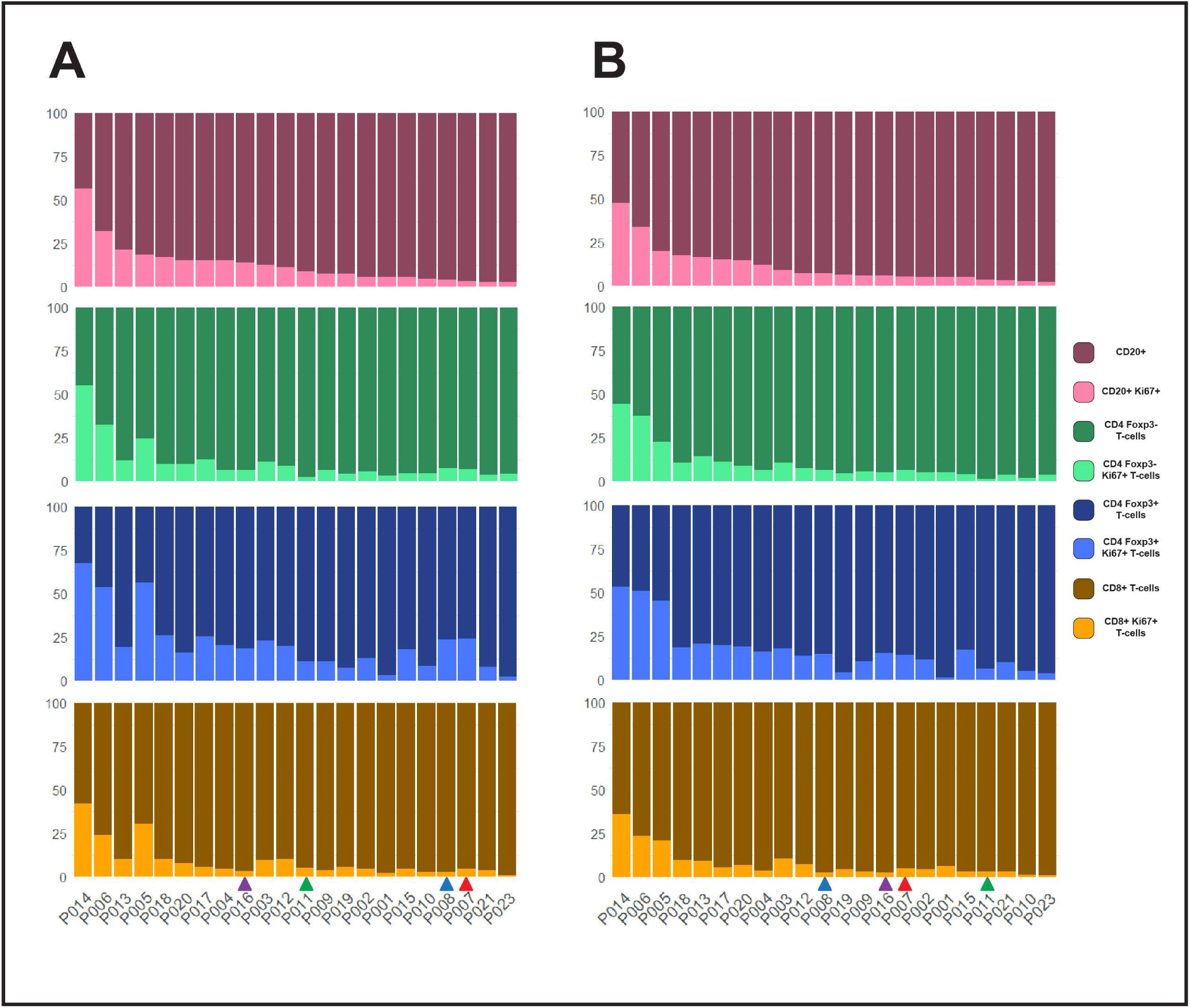
Rankings of patients by quantification of B-cell Ki-67 in individual TLS domains vs all TLS domains combined. 4 lymphocyte subsets were identified within TLS domains and the proportions of Ki-67+ cells within those subsets were calculated and plotted in stacked plots. Patients were ranked according to the proportion of B-cells expressing Ki-67, calculated according to two different methods. **(A)** As for Figure 3B and 3C, these proportions were calculated for each individual TLS domain then averaged. **(B)** As for Figure 6A, cell phenotypes were quantified across all pooled TLS domains within the tissue sample. Coloured arrows are used to highlight four patients (P007=red, P008=blue, P011=green, P016=purple) who had significant changes in rank between the two different approaches.

**Figure S3:**
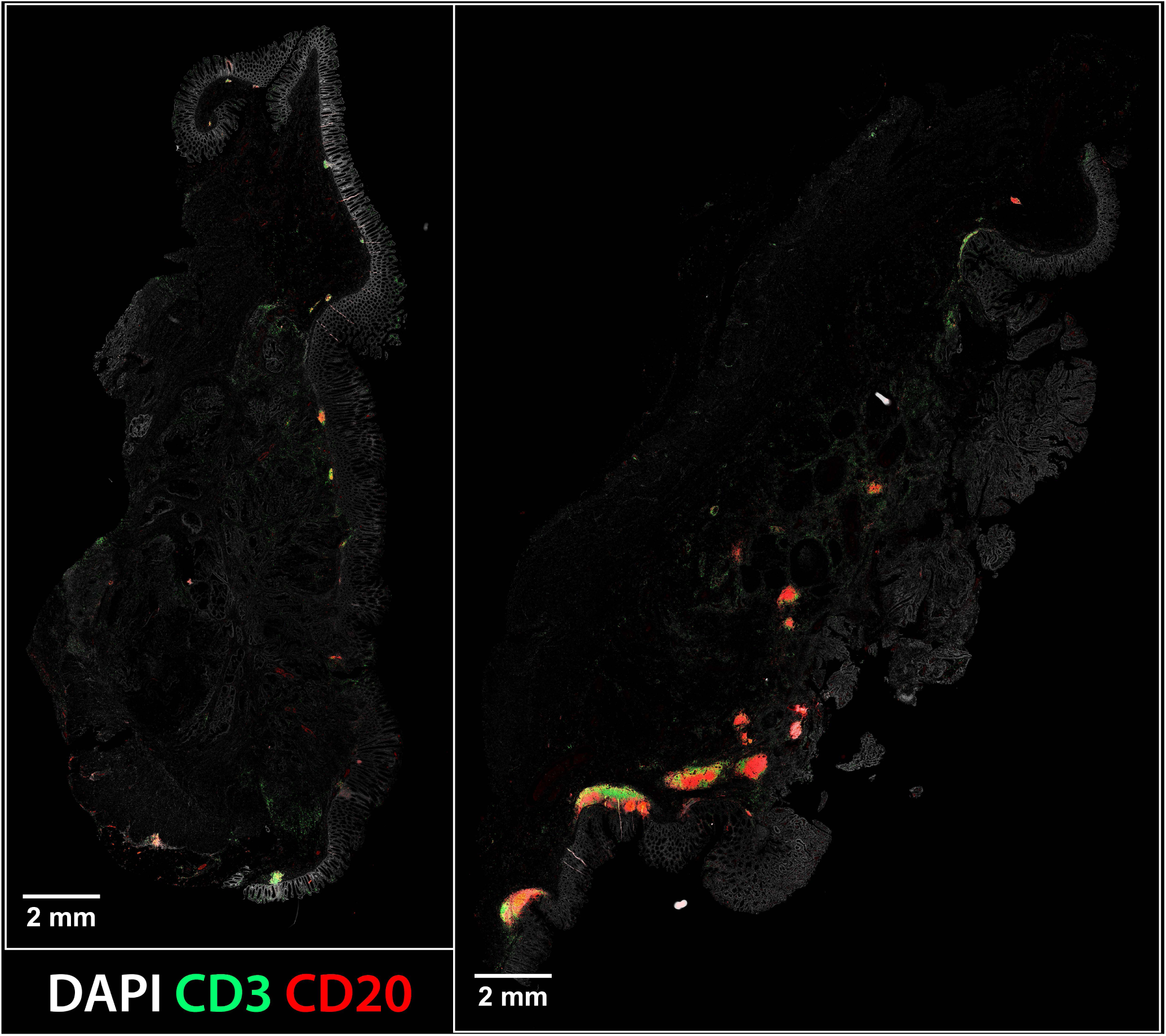
Disparity between the count of TLS domains and total area of TLS. Whole scan images of tissue sections from P006 (left) and P013 (right), showing only DAPI (light grey), CD3 (green) and CD20 (red).

